# msengine: Adaptive Open-Source Signal Extraction for HDX MS

**DOI:** 10.64898/2026.09.30.755802

**Authors:** Alexander Patapoff, Nicholas Patapoff, Benjamin Walters

**Affiliations:** Departments of Biochemistry and Cellular Pharmacology, Genentech, Inc.; Departments of Biochemistry and Cellular Pharmacology and Structural Biology, Genentech, Inc.

## Abstract

Hydrogen-deuterium exchange mass spectrometry (HDX-MS) requires extraction of broad, time-dependent peptide isotope envelopes from noise and overlapping signals. The ExMS/ExMS2 lineage established a retention-time-aware framework for raw-data extraction and integrated HDX-MS analysis. Building on this foundation, msengine is an open-source Python engine that adapts mass-selection windows and retention-time boundaries to the observed data. Spectra are reconstructed as nonnegative combinations of the theoretical undeuterated (all-H) isotope distribution shifted across allowed deuteration states, without prescribing a population shape. The fitted weights, termed deuteration coefficients, describe each state’s contribution and guide refinement of mass-selection windows. Spectral matches establish an initial retention interval, which is then expanded using chromatographic intensity to include weaker portions of the elution profile. Interference screening and final quality checks assess the recovered interval. In a 1:1 mixture of undeuterated and deuterated material, interval expansion changed the relative population intensities within the extracted envelope resulting in closer agreement to the 1:1 target, illustrating the importance of boundary selection. Across a four-timepoint experiment with the SET1/SWD2 complex collected in duplicate, msengine accepted an average of 901 peptide–charge targets per measurement versus 636 for ExMS2, a 41.7% increase, while improving replicate-centroid disagreement and retention-window widths between replicates. In a same-workstation benchmark, total wall time decreased from 71.9 to 7.3 min, with higher peak memory use. Together, these results demonstrate adaptive signal extraction across mass and retention time as a means to retain multimodal envelopes while increasing accepted-target yield, persistence, accuracy, and throughput.

## Introduction

Hydrogen-deuterium exchange mass spectrometry (HDX-MS) measures protein conformational dynamics, interactions, and structural heterogeneity in solution, and is widely used to localize binding interfaces, allosteric changes, folding events, and protein-state differences.^1,2^ In the common bottom-up workflow, this information is distributed across many partially deuterated peptides, making raw-signal extraction a prerequisite for quantitative interpretation. Peptide identifications and undeuterated reference spectra define the extraction targets, while deuterated LC-MS runs produce broad, time-dependent isotope envelopes that must be isolated reproducibly. This step is foundational: centroids, uptake values, population fits, differential maps, and structural interpretations all inherit the quality of the extracted signal.

Substantial software development has advanced HDX-MS analysis, particularly for visualization, statistical comparison, multimodal analysis, back-exchange correction, and higher-resolution interpretation after peptide envelopes have been assigned.^2^ By comparison, fewer tools focus on upstream raw-data extraction itself, even though this is where HDX-MS data first become quantitative. ExMS2,^3^ building on ExMS and HDsite,^4,5^ is an important contribution in this space, providing an integrated workflow for peptide finding, envelope recording, assignment validation, multimodal analysis, and higher-resolution interpretation. HeXicon and Hexicon 2^6^ similarly recover full deuteration distributions from theoretical isotope representations; msengine uses this representation as a scan-level consistency test within an ExMS2-style retention-time-aware workflow. Nevertheless, assigning deuterated envelopes from LC-MS data remains difficult to automate with minimal parameters and without reliance on downstream physical models. Extraction must reject chemical noise, contaminants, coeluting peptides, carryover, retention-time outliers, and overlapping charge states while preserving the target peptide’s full chromatographic and isotopic signal—or flagging the measurement when that signal cannot be isolated reliably. Existing workflows therefore often use heuristics about centroid behavior, envelope continuity, retention-time consistency, or deuterium-distribution shape to reduce false assignments. These criteria help suppress false assignments, but when used to define signal membership they can favor expected envelope shapes and under-represent heterogeneous or multimodal exchange; relaxing them without an alternative strategy risks absorbing overlapping or aliased signal into the target measurement. These competing requirements motivate an extraction strategy that rejects interference by physical inconsistency with the target ion, rather than by enforcing a predefined deuterium-distribution shape.

Here we present msengine, an open-source, lightweight, Pythonic, extensible, and multithreaded engine for extracting deuterated isotope envelopes from HDX-MS data. msengine supports standard fragment-separation HDX-MS workflows and can also perform targeted extraction of user-specified deuterated ions from a chemical formula and exchangeable-site count, making it applicable beyond canonical peptide targets. Continuing the retention-time-aware raw-data philosophy of ExMS2, ^3^ msengine decouples signal extraction from distributional modeling while minimizing user-supplied processing parameters. Envelope membership is defined by exact theoretical isotopologue positions, and arbitrary deuterated distributions are reconstructed by non-negative forward substitution of the theoretical all-H distribution across allowed deuteration states. The resulting deuteration-vector representation asks whether the observed signal can be physically constructed from the target molecule, rather than whether it conforms to a binomial, Gaussian, or smoothly evolving envelope. To maximize data fidelity without manual intervention or extensive user-tuned heuristics, msengine uses instrument PPM error as the principal user-supplied parameter and derives resolving power, mass-bin refinement, and retention boundaries from the measured spectra and chromatographic signal. Resolving power is estimated from each reference spectrum; isotopologue-aware extraction bins are constructed from theoretical peak positions and tightened after an initial deuteration vector is solved; a retention interval is first identified from per-scan spectral agreement (the seed interval) and then expanded using chromatographic intensity to include weaker portions of the elution profile; and higher-charge-state aliasing is detected, used to refine the retention window, and incorporated into quality-control criteria that determine whether the interval is retained for downstream processing. Because each processing stage is implemented as a modular Python component, the workflow can be run end-to-end or customized for specialized extraction, filtering, or downstream modeling. By preserving complex deuterated envelopes first and modeling them later, msengine provides an automated, scalable extraction layer for modern HDX-MS workflows and instrumentation.

## Methods

### Pipeline Outline

The msengine pipeline executes in eight stages, numbered 0 through 7, beginning with the experimental data collection that defines the input and proceeding through the computational steps that produce per-peptide deuteration distributions with associated quality metrics.

#### Stage 0: Experimental data collection

The standard hydrogen-deuterium exchange mass spectrometry (HDX-MS) workflow comprises three phases:^1^ peptide identification, an undeuterated (“all-H”) reference measurement, and measurement of time-dependent deuterium uptake. The outputs of these phases (a peptide list with charge states, the all-H reference spectrum, and the time-resolved deuterated spectra) constitute the input to the computational pipeline.

#### Stage 1: Reference ingestion and bin construction

The engine inputs the peptide list, charge states, raw all-H reference spectrum, and the instrument’s ppm mass accuracy. For each peptide, the resolving power *R* is determined empirically from the spread of the monoisotopic peak in the all-H spectrum. The peptide-specific *R* and ppm tolerance are used to construct an initial set of static bins covering every theoretically possible deuterated peak position for the target envelope (see *Segmented-Scan Spectral Stitching: Static Binning*).

#### Stage 2: Per-scan peak centroiding

For each scan, the raw profile data spanning the peptide’s *m/z* range (extended by one charge-spacing unit on each side to capture peaks adjacent to the envelope) are reduced to a set of (*m/z*_center_, *I*_integrated_) peak measurements using the Partial Area Gaussian Estimation (PAGE) method (see *Multimodal Peak Determination: Shape-Ratio Analysis*). PAGE fits the local profile shape of each peak and integrates intensity beneath it, producing a sparse list of integrated peak measurements across the peptide window.

#### Stage 3: Envelope construction and deconvolution

The peak measurements from each scan are assigned to the static bins by *m/z* position, and the integrated intensities of all measurements falling within a given bin are summed to produce that bin’s entry in the extracted-ion-chromatogram (XIC) mass-envelope vector. The envelope is deconvolved by constrained forward substitution (see *Envelope Validation*), producing a per-scan deuteration distribution and an associated *R*^2^ goodness-of-fit metric.

#### Stage 4: Seed interval selection

A seed retention-time interval is identified from the contiguous scans whose *R*^2^ values exceed a user-specified threshold. Candidate intervals are scored by duration, mean fit, and mean intensity, and the highest-scoring candidate is validated against the run-wide distribution of apex deviations established from the undeuterated reference pool; outliers are replaced by the highest-scoring non-outlier candidate (see *Initial Seed Interval*). The per-scan XIC envelopes within the surviving interval are aggregated into a single seed-level envelope that anchors the recalibration of Stage 5.

#### Stage 5: ppm recalibration and interval extension

The seed interval anchors the procedure described under *Dynamic ppm Recalibration*, which tightens the static bins to the subset of theoretical positions consistent with the deconvolved deuteration. The retention interval is then reprocessed and extended outward from the seed to capture the full chromatographic peak.

#### Stage 6: Aliasing exclusion

The extended retention window is screened for interference from overlapping higher-charge species using the cosine-squared mask described under *Overlapping Charge State Interference: Anti-Aliasing*. Scans whose aliased contribution exceeds the threshold criterion are excluded, and the retention interval is contracted accordingly.

#### Stage 7: Final review and flag consolidation

After the per-peptide pipeline completes, each peptide’s accumulated state is reviewed against the criteria defined under *Quality Control Flagging*. Flags raised tentatively during earlier stages are retained or cleared based on this review, and any conditions detectable only from the finalized state are flagged at this point. The resulting flag set is emitted with the peptide’s other outputs.

### Multimodal Peak Determination: Shape-Ratio Analysis

In mass spectrometry, distinct molecules or isotopes with similar *m/z* values often produce underlying signals that merge into a single, structurally unresolved cluster of peaks in the raw data.^7–10^ To resolve these clustered raw data points when feasible, we implemented a hybrid decomposition strategy using Gaussian kernels; substantial overlap (e.g., |*µ*_1_ *− µ*_2_|*≤ σ*_1_ +*σ*_2_) prevents decomposition. Initial parameters for the underlying signals are rapidly estimated using Partial Area Gaussian Estimation (PAGE), described in the Supporting Information. Composite peaks are deconvolved by iteratively expanding the model. A sequential Bayesian information criterion (BIC) determines when to stop adding Gaussian components, while the Runnalls reduction algorithm^11^ evaluates whether the final model complexity should subsequently be reduced. Figure 1 illustrates the method’s performance for three overlapping peaks, one of which is significantly obscured by its neighbors at a 10:1 signal-to-noise ratio (SNR).

**Figure 1.**
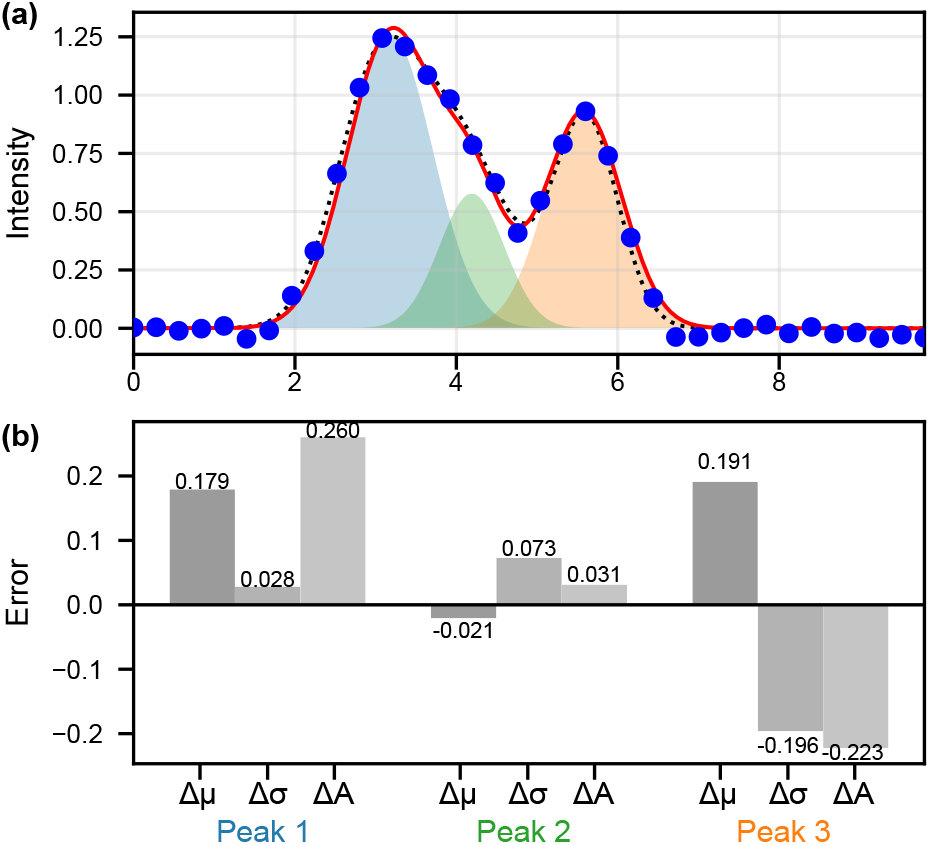
PAGE decomposition of three overlapping Gaussians at low resolution (~2.5 points/*σ*). **(a)** Blue circles, noisy data; black dotted line, true composite signal; colored fills, individual PAGE-fitted Gaussian components; red solid line, total PAGE fit. **(b)** Per-peak parameter errors (fitted minus true). Blue bars, position error (Δ*µ*); orange bars, width error (Δ*σ*); green bars, amplitude error (Δ*A*).

### Segmented-Scan Spectral Stitching: Static Binning

A static binning procedure establishes the set of *m/z* intervals (bins) where peaks from a target mass envelope might reside, and it must accommodate every relevant isotopologue (using an abundance threshold of 0.1%) across all possible deuteration states. First, the natural-abundance (all-H) isotopic distribution of the target peptide is computed, yielding a set of *m/z* peak centers and intensities (Figure 2a). This distribution is then translated through every accessible deuteration state from +0 to +*N* uptake sites, with each shifted copy contributing its *m/z* peak centers to a unioned theoretical envelope (Figure 2d shows the first union). From these coordinates, bins are constructed in a two-pass procedure that accounts for the finite resolving power of the instrument, which sets the minimum distinguishable separation between *m/z* peak centers, and ppm error, which broadens the acceptable range around each theoretical position. The resulting bins span every *m/z* position the peptide can produce under any sequence-consistent deuteration configuration while remaining as small as possible to eliminate interference from peaks that are not part of the target envelope.

**Figure 2.**
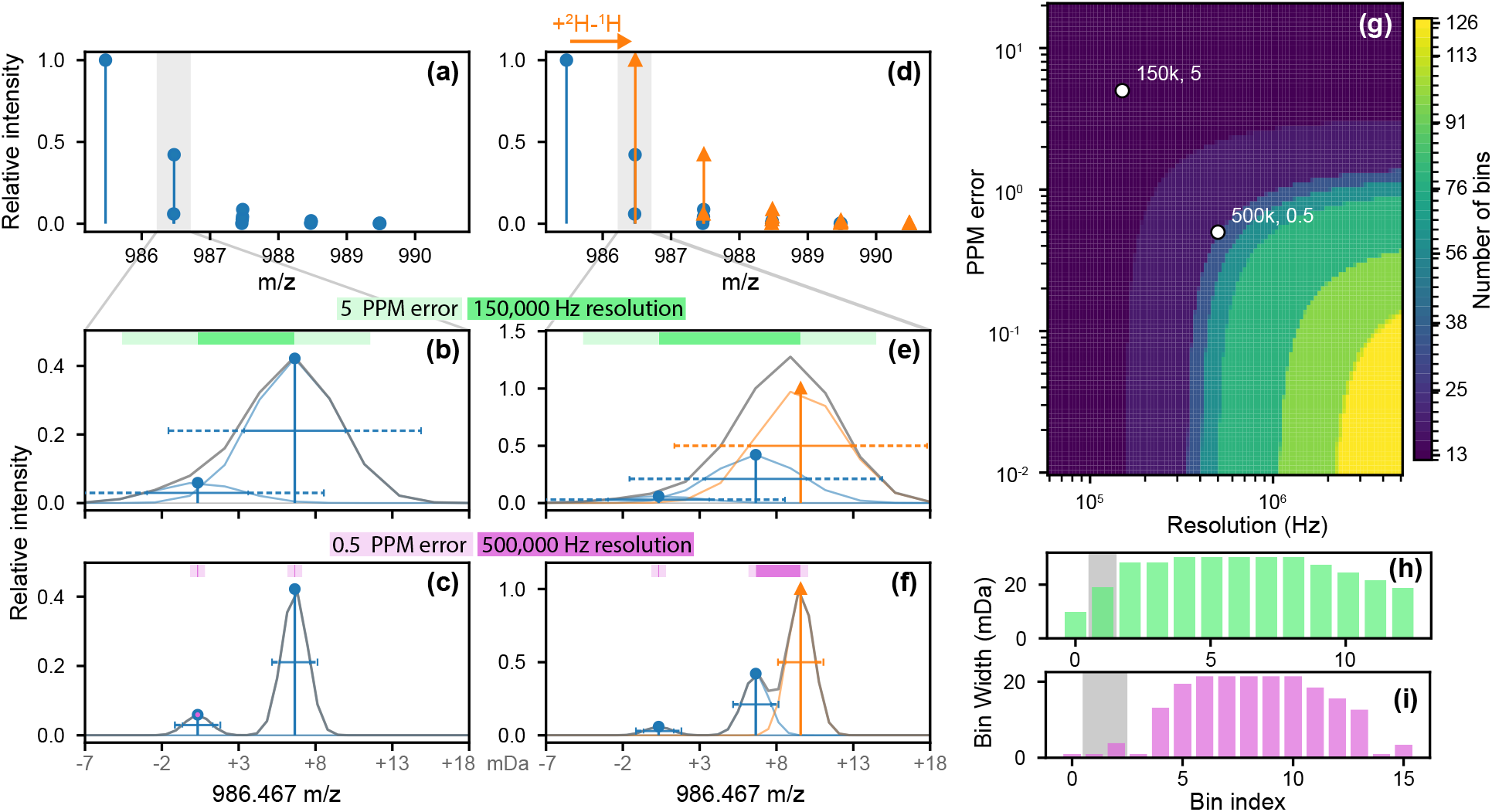
Centroid-resolution and mass-drift binning illustrated on the +1 charge-state isotopic envelope of the peptide HDISGREAT (monoisotopic *m/z ≈*986.46). Throughout, blue circles denote the all-H isotopic distribution and orange triangles denote the same distribution shifted right by one deuteron mass. **(a)** Theoretical all-H stick spectrum across the full envelope. **(d)** Combined all-H +1D stick spectrum across the full envelope. **(b, c)** Simulated response at the +1 nominal isotope bin for the all-H distribution in (a), at two regimes: **(b)** *R* = 150,000, ppm error = 5.0 (loose, low-resolution) and **(c)** *R* = 500,000, ppm error = 0.5 (tight, high-resolution). **(e, f)** Simulated response at the +1 nominal bin for the all-H +1D distribution in (d), at the same two (*R*, ppm error) settings. In the zoom panels, the gray curve is the sum of all per-centroid Gaussians broadened to FWHM = *m/R*; per-stick horizontal whiskers at *y* = *I/*2 show each centroid’s FWHM*/*2 (solid) and *±* ppm error extension (dashed). The colored bar near the top of each panel marks the resulting final bins: the inner solid segment spans the lowest-to-highest member *m/z* of that bin, the lighter shaded shoulders mark the additional *±* (ppm error*/*10^6^) *· c* tolerance, and the white diamond marks the bin’s intensity-weighted center. **(g)** Heatmap of the number of final bins produced by sweeping *R ∈* [6 *×* 10^4^, 5 *×* 10^6^] against ppm error *∈* [10^*−*2^, 20] over the full deuterated stick spectrum (all uptake states 0, …, *n*_D_). As the two sources of uncertainty shrink (high *R*, low ppm error; lower-right of the map), more peaks resolve into separate bins; as either tolerance grows (upper-left), centroids merge and the bin count collapses. White circles labeled “row 1” and “row 2” mark the parameter values used for rows (b, e) and (c, f), respectively. **(h, i)** Per-bin width in mDa for those two settings (**h**: row-1 parameters, coral; **i**: row-2 parameters, steel-blue), showing the width distribution across the final bins.

#### First pass: resolution-based grouping

For each theoretical *m/z* peak center at position *x*_*i*_, a resolution-dependent half-width is computed from the instrument’s resolving power *R*,

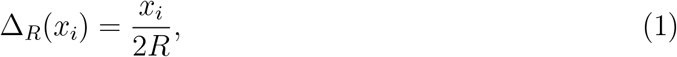

defining the interval [*x*_*i*_ *−*Δ_*R*_(*x*_*i*_), *x*_*i*_ + Δ_*R*_(*x*_*i*_)] whose total span equals the instrument’s full width at half maximum at that *m/z. m/z* peak centers whose intervals overlap cannot be resolved by the instrument and are merged into a single group by traversing the sorted intervals from low to high *m/z*. Merging continues across consecutive overlaps until a clear gap is encountered causing the group to close. Each closed group yields a preliminary bin whose center is the arithmetic mean of the contributing *m/z* peak centers and whose intensity is the sum of their intensities. The minimum and maximum *m/z* peak center positions (*x*_*min*_, *x*_*max*_) within the group are retained for use in the second pass.

#### Second pass: ppm expansion and consolidation

These preliminary bins are then expanded to absorb scan-to-scan variation (ppm error). For a preliminary bin with center 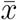, a ppm half-width is computed as

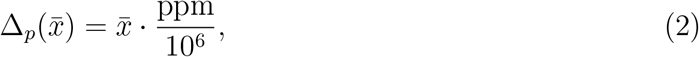

and the bin’s first-pass interval edges are widened by 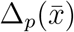 on each side. The expanded intervals are traversed a second time in *m/z* order, and any preliminary bins whose expanded intervals overlap are consolidated into a single final bin. The center of each final bin is the midpoint of its smallest and largest contributing *m/z* peak center positions,

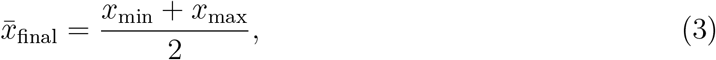

and its width is the span between those positions, padded by 2Δ_*p*_ evaluated at that center,

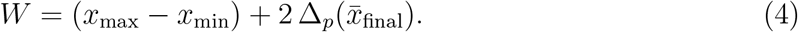

This yields the static bin configuration: a set of disjoint *m/z* intervals that accommodate every theoretical isotopologue position the peptide can produce across all deuteration states, with bin geometry scaled to the instrument’s resolving power and mass accuracy (Figure 2b– i).

### Envelope Validation

Arbitrary deuterated distributions can be reconstructed from a deuteration-vector representation 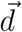, where each element *d*_*i*_ is a non-negative coefficient representing the fraction of the population that has incorporated exactly *i* deuterons. The observed binned intensity vector 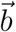 is then expressed as the convolution of 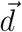 with the theoretical all-H isotopic distribution, which can be written as the linear system

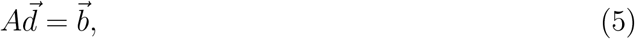

where *A* is a lower-triangular matrix whose columns are successive integer-mass shifts of the binned all-H envelope. The matrix is lower-triangular because deuteration only displaces signal toward higher mass: a sub-population with *i* deuterons contributes to bin *j* only when *j ≥ i*.

The system is solved by constrained forward substitution, building on prior deconvolution approaches in HDX-MS.^12,13^ Because *A* is lower-triangular, the coefficients *d*_*i*_ can be computed sequentially from *i* = 0 upward, with each step requiring only the already-determined coefficients *d*_0_, …, *d*_*i−*1_ and the observed intensity *b*_*i*_. At each step, forward substitution yields a candidate coefficient algebraically; the algorithm accepts the candidate only if it is non-negative. Steps that violate this condition are rejected and the corresponding coefficient is set to zero, since a negative coefficient would correspond to a nonphysical population.

In intuitive terms, the algorithm deconvolves the spectrum iteratively: at each step it scales the theoretical all-H envelope by the candidate coefficient, subtracts this contribution from the residual observed signal, and advances by one exchange unit to resolve the next coefficient. Figure 3a illustrates this on a composite envelope constructed from four exchange populations, including a well-separated sub-population shifted four bins from the main distribution; the reconstruction recovers the deuterium uptake distribution across the full envelope from the observed signal alone.

**Figure 3.**
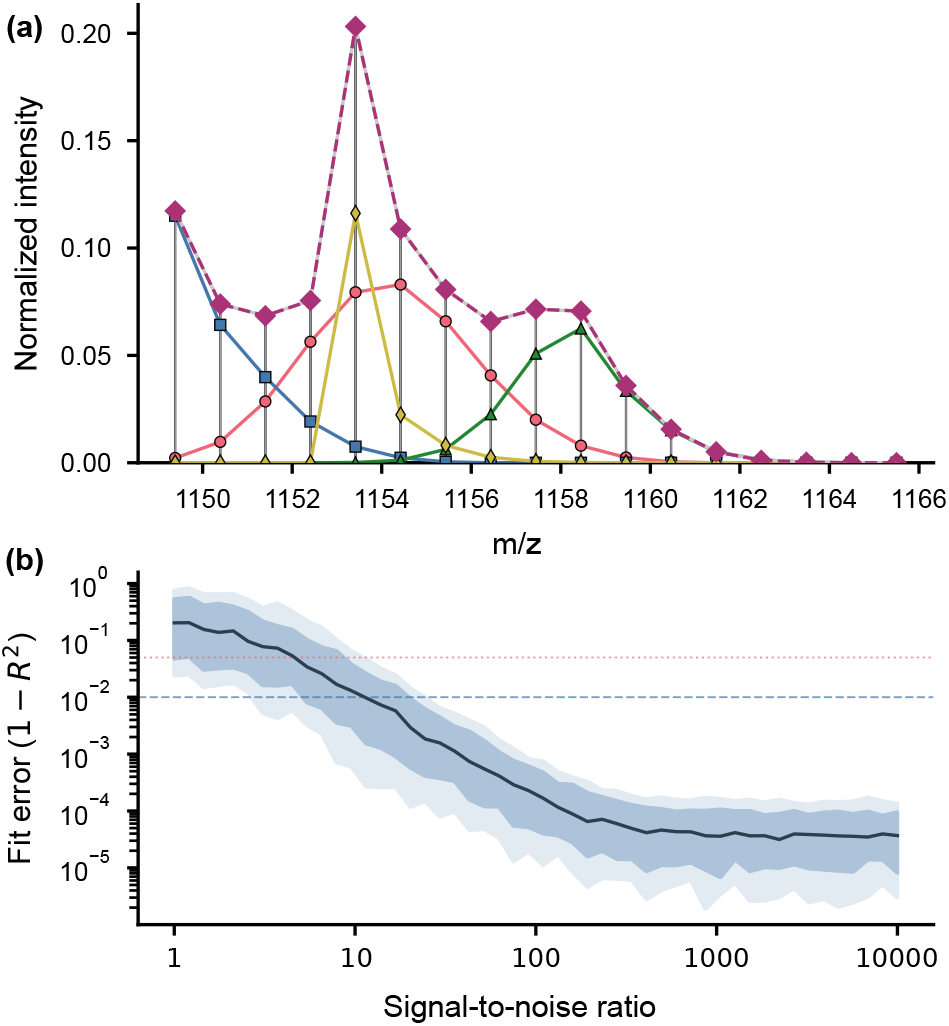
Isotopic envelope validation using modified forward substitution. **(a)** Simulated composite isotopic envelope (gray bars and line) constructed from four deuterium-exchange populations: A (*f*(*D*) = 0.5, 40%, coral), B (*f*(*D*) = 0.1, 25%, blue), C (*f*(*D*) = 0.9, 20%, green), and D (*f*(*D*) = 0.4, four sites; *f*(*D*) = 1 for the remaining sites, 15%, yellow), where *f*(*D*) represents fractional deuterium carried. The best-fit reconstruction (purple diamonds, dashed) overlays the input mixture. **(b)** Fit error (1*− R*^2^) as a function of additive Gaussian noise level, expressed as a fraction of maximum signal intensity. The dark line shows the median across 200 replicates per noise level; inner and outer shaded bands represent the *±*1*σ* and *±*2*σ* intervals, respectively. Horizontal lines mark *R*^2^ = 0.99 (blue, dashed) and *R*^2^ = 0.95 (coral, dotted) thresholds.

### Initial Seed Interval

The seed interval is the smallest, highest-confidence stretch of scans within the peptide’s expected elution window, identified to anchor the dynamic ppm recalibration before the retention interval is extended.

A search window centered on the reference retention time is established, with a half-width of Δ*t*_H_ for all-H reference data and Δ*t*_D_ for deuterated samples. Within this window, contiguous scans are identified for which the per-scan isotopic goodness-of-fit metric *R*^2^ defined under *Envelope Validation* exceeds a user-specified threshold (default 0.7). A one-scan look-ahead bridges single-scan failures within an otherwise contiguous run, accommodating transient noise.

A peptide’s true elution event can be fragmented into several disconnected intervals by noise or by interference from overlapping envelopes, so all candidate intervals are scored and ranked. Each is assigned a score

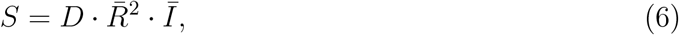

where *D* is the interval duration in number of scans, 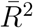 is the mean goodness-of-fit, and *Ī* is the mean integrated intensity. The highest-scoring interval is selected as the tentative seed. If no interval clears the *R*^2^ threshold, the spectra are smoothed with a moving-average filter spanning approximately half the expected elution width and the search is repeated; this fallback recovers low-SNR peaks whose per-scan fits are individually weak but collectively coherent.

Because a high local score does not guarantee a correct assignment, the tentative seed is then checked against the run-wide elution behavior. For each peptide, the deviation between its tentative apex and its reference apex is computed; the set of all such deviations forms an empirical distribution of retention-time shift across the run, established from the undeuterated reference pool.^1^ Outliers are identified by the interquartile range (IQR) of this distribution: tentative values whose deviations fall outside the two-sided fence [*Q*_1_ *–* 1.5 IQR, *Q*_3_ + 1.5 IQR] are rejected. When the highest-scoring candidate is rejected, the next-highest is tested against the same fence, and the procedure continues down the ranked candidates until one falls within bounds; if none qualifies, the lowest-deviation candidate is retained. This catches cases where the highest-scoring interval is locally compelling but inconsistent with the run’s overall chromatography. The resulting seed anchors the procedure described under *Dynamic ppm Recalibration*.

### Retention Interval Consistency Check

For deuterated runs, retention-time outlier rejection is performed by comparing each peptide’s measured retention interval against its all-H reference interval after correcting for retention-time drift between the two runs. This replaces the all-H apex-deviation check described under *Initial Seed Interval* and is applied to all peptides except those flagged as bad data or processing errors under *Quality Control Flagging*.

#### Drift correction

For each peptide *p*, the apex offset between the deuterated and all-H runs is

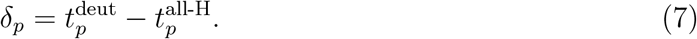

A global drift estimate *δ*_global_ is taken as the median of *δ*_*p*_ across all peptides. To capture drift that varies across the gradient, peptides are also grouped by their all-H apex into bins of width Δ*t*_bin_ (default 0.5 min), and a local drift estimate *δ*_local_(*t*) is taken as the median of *δ*_*p*_ within each bin. For a given peptide, the local estimate is used when its bin contains at least *N*_min_ peptides (default 10); otherwise the global estimate is used.

#### Consistency check

The all-H reference interval [*a*_H_, *b*_H_] is shifted by the applicable drift estimate *δ* to produce the expected deuterated interval *E* = [*a*_H_ + *δ, b*_H_ + *δ*], which is compared against the measured deuterated interval *I* = [*a*_D_, *b*_D_] by intersection-over-union (Jaccard index),

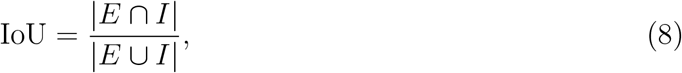

the length of the intervals’ overlap divided by the length of their union, taken as 0 when they are disjoint. Peptides with IoU *≤ τ*_IoU_ (default 0.25) are flagged as RT-inconsistent.

### Dynamic ppm Recalibration

Following initial static calibration and data extraction over the seed interval, a dynamic ppm recalibration step refines the bin configuration using the deuterium distribution from the preceding deconvolution. The static bins, constructed to accommodate the union of all theoretically possible deuterated isotopologues, are necessarily broad because the real distribution rarely populates every accessible deuteration state. Recalibration narrows these bins to the subset of theoretical positions supported by the deconvolved coefficients described under *Envelope Validation*.

The refined theoretical distribution is constructed by convolving the deconvolved deuteration coefficients 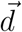 with the peptide’s all-H isotopic distribution. This yields a list of theoretical peaks indexed by *i*, where each peak *i* has an *m/z* position *x*_*i*_ and an intensity *w*_*i*_ representing its contribution to the overall envelope under the deconvolved distribution. Peaks with *w*_*i*_ = 0 are dropped; the remaining set *{* (*x*_*i*_, *w*_*i*_) *}* contains only the theoretical positions actually supported by the data.

These peaks are then used to create new bins. The list is sorted by *x*_*i*_ and traversed from low to high *m/z*; a new bin opens whenever the gap to the preceding peak exceeds twice the user-defined ppm uncertainty,

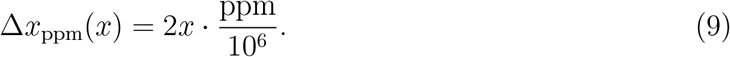

Within an open bin *B*, the refined center is the intensity-weighted mean of the contributing peak positions,

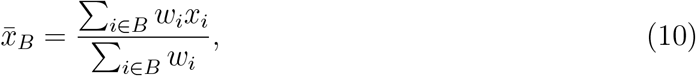

and the intensity-weighted variance of those positions is

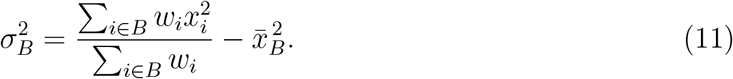

The bin width is then defined as

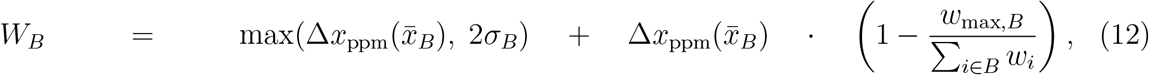

where *w*_max,*B*_ = max_*i∈B*_ *w*_*i*_ is the largest peak intensity within the bin and the index *i* ranges over peaks of the refined distribution that fall within *B*.

The first term enforces a floor at the ppm-derived measurement uncertainty, ensuring no bin is narrower than the calibrated error of the instrument. The second term inflates the bin in proportion to how diffuse the solved intensity is across contributing peaks: when a single peak dominates 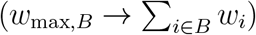, the inflation vanishes and the bin remains at its measurement-floor width; when intensity is spread evenly across multiple peaks, the inflation approaches a full ppm window, accommodating the ambiguity in the underlying distribution. All spectral data are subsequently re-extracted against this refined bin configuration.

### Retention Interval Expansion

Once the seed interval [*a*_0_, *a*_1_] is established and the bins are recalibrated as described under *Dynamic ppm Recalibration*, the interval is extended outward to capture the full chromatographic peak. The per-scan *R*^2^ threshold alone would truncate the profile at its low-intensity edges, where the fit degrades although the signal is genuine, so the expansion terminates instead on observable chromatographic features.

Let *I*_*i*_ be the summed intensity at scan *i* of the most abundant isotope peaks, those whose matched relative abundance lies within 10% of the envelope maximum. Single-scan spikes, which continuous elution cannot produce, are first replaced by their taller neighbour wherever they exceed it threefold. Three quantities are then computed and held fixed. The flank *ℱ* holds the scans more than *p* from either seed edge. The floor *β* is the median of its non-zero scans. The ceiling *c* = med *ℱ* + *z* max 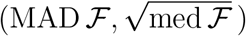 uses every flank scan, zeros included, with MAD the median absolute deviation scaled to estimate a normal standard deviation. The third quantity, the relative noise *ν*, derives from the second differences *d*_*j*_ of the positive intensities near the seed, normalised to unit variance. This is insensitive to local level and slope, and reflect measurement error rather than peak shape, giving *ν* = MAD(*d*)*/L* clipped to [0.02, 1] with *L* their median.

Each side walks outward from its seed edge, independently of the other, tracking the largest intensity seen so far, *M*, the smallest seen since that maximum, *m*, and the scan *i*_*m*_ where *m* occurred. A scan at or below *β* updates none of them, so a dropout is not mistaken for a valley. A new maximum resets all three, so depth is always measured from the most recent peak. Two criteria are then evaluated. The first applies where *I*_*i*_ *> β*, terminating expansion at *i*_*m*_ if

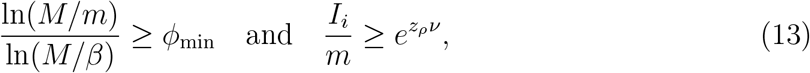

the trace having descended through a fraction *ϕ*_min_ = 0.3 of its logarithmic span and since recovered by more than noise admits (*z*_*ρ*_ = 20). The second applies at every scan: *I*_*i*_ *< θ* sustained over *n*_*c*_ scans ends the walk at the last scan that passed, where *θ* = max(*βe*^*zν*^, *c*) and *z* = 2. Requiring a run prevents an isolated excursion from truncating the peak, and scans at or below *β* still count toward it. Reaching the end of the acquisition terminates the walk at the same edge. parameter values were selected against a sweep of synthetic noisy background models.

### Overlapping Charge State Interference: Anti-Aliasing

Each scan’s expected mass envelope is evaluated to determine whether it corresponds to a higher charge state with similar mass at the given resolution and ppm error of the data. This involves the application of a dual-thresholded cosine-squared mask, with the origin shifted to the expected monoisotopic mass of the target peptide. The method acts as a matched filter, probing the spectrum for specific isotopic periodicities relative to the expected distribution while continuously accounting for calibrated measurement error. Two parallel conditions account for the distinct isotopic mass spacings of carbon (Δ*M*_*C*_ = ^13^C *−* ^12^C) and hydrogen (Δ*M*_*D*_ = ^2^H *−* ^1^H). Rather than using static mass windows, the acceptable mass tolerance for a peak center is mathematically dictated by the amplitude thresholds (*y*_thresh,*D*_, *y*_thresh,*C*_) applied to these periodic masks, where higher thresholds correspond to narrower allowed windows. By adjusting these thresholds, the allowable *m/z* width around each theoretical peak position expands or contracts proportionally to the ppm error, eliminating signals that fall outside these boundaries and enabling the algorithm to function dynamically at any resolution.

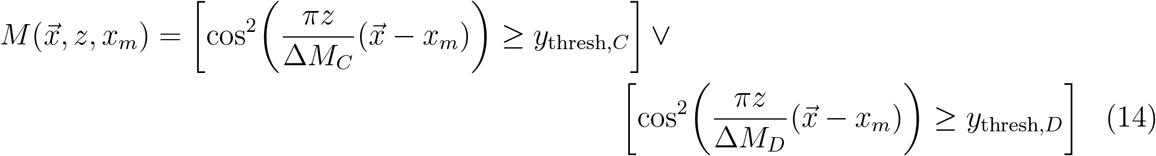

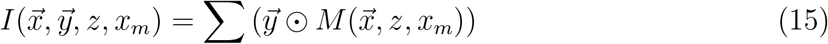

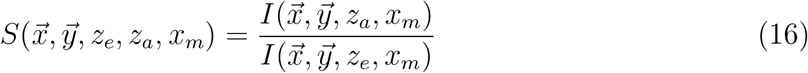

Here, 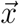 and 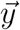 are the vectors of *m/z* peak positions and intensities, and *x*_*m*_ is the shifted monoisotopic origin. *M* 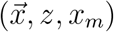 is the boolean mask evaluating the charge state *z* against the expected isotopic mass differences Δ*M*_*C*_ and Δ*M*_*D*_, controlled by the thresholds *y*_thresh,*C*_ and *y*_thresh,*D*_ respectively; the mask is treated as a 0/1 indicator vector when combined with 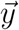 via the Hadamard product. *I* is the masked intensity sum at a given charge state. The score *S* compares the intensity captured at an alias charge *z*_*a*_ to that captured at the expected charge *z*_*e*_, where *z*_*e*_ is fixed at the peptide’s nominal charge state while *z*_*a*_ is swept across higher candidate charges. Increasing *z*_*a*_ raises the mask frequency, allowing the integration to capture intensity at *m/z* positions consistent with higher-charge aliased species. The masks at *z*_*a*_ and *z*_*e*_ probe distinct but partially overlapping *m/z* positions, so *S* reflects the relative intensity contribution attributable to a higher-frequency alias without requiring one mask to subsume the other.

As demonstrated in Figure 5, the method probes the spectrum for specific isotopic periodicities relative to the expected distribution while continuously accounting for calibrated measurement error. To systematically detect and flag aliased interference, a hard cutoff threshold of *S ≥* 1.5 is implemented. This operating point indicates that intensity captured at the alias charge exceeds the expected-charge intensity by 50% or more, prompting the algorithm to exclude the contaminated signal.

**Figure 4.**
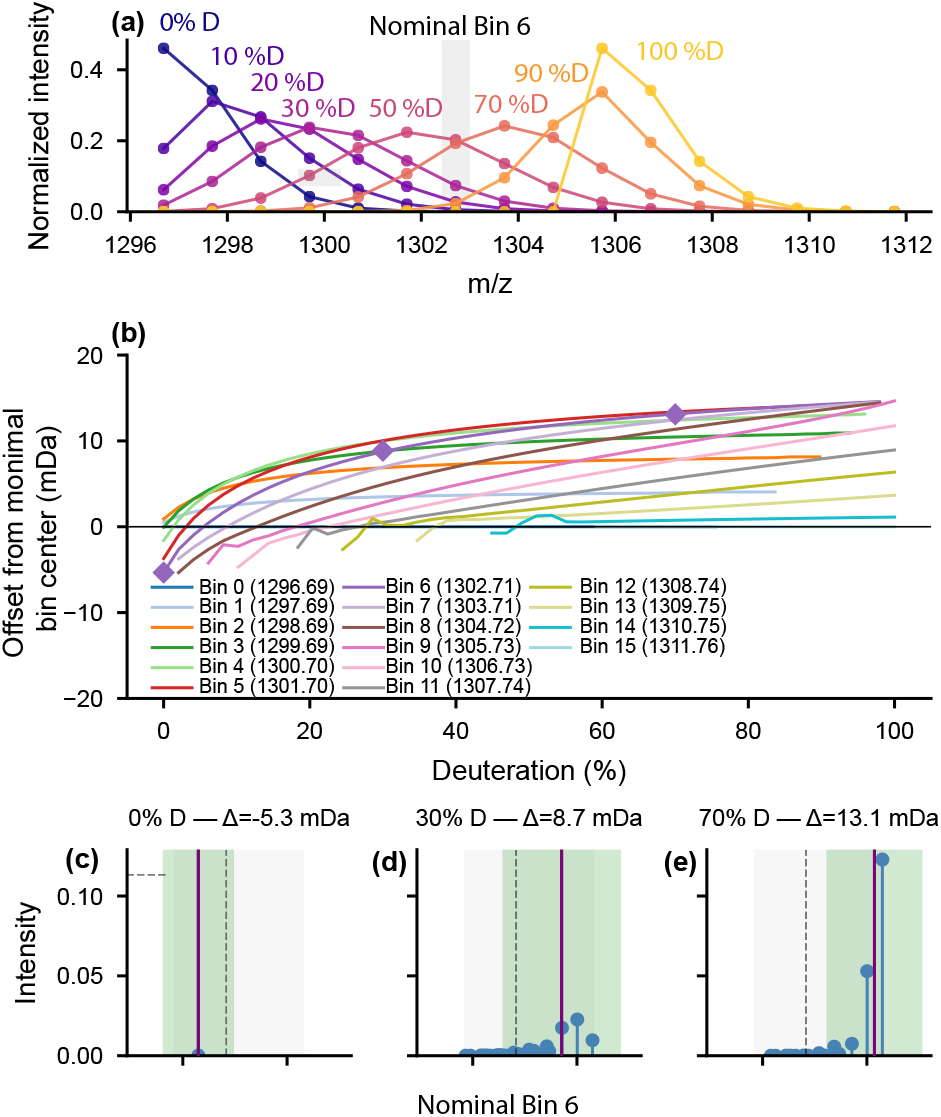
Deuteration-dependent mass drift within nominal bins. Simulations used a singly charged peptide (DRVYIHPFHL). **(a)** Binned isotopic envelopes across 0–100% deuteration; gray shading marks bin 6. **(b)** Centroid offsets from nominal bin centers as a function of deuteration; diamonds mark the conditions expanded below. **(c–e)** Bin 6 at 0%, 30%, and 70% deuteration. Blue sticks are isotopic peaks, the dashed black line is the nominal center, green regions and lines are recalibrated bins, and the purple line is the intensity-weighted centroid.

**Figure 5.**
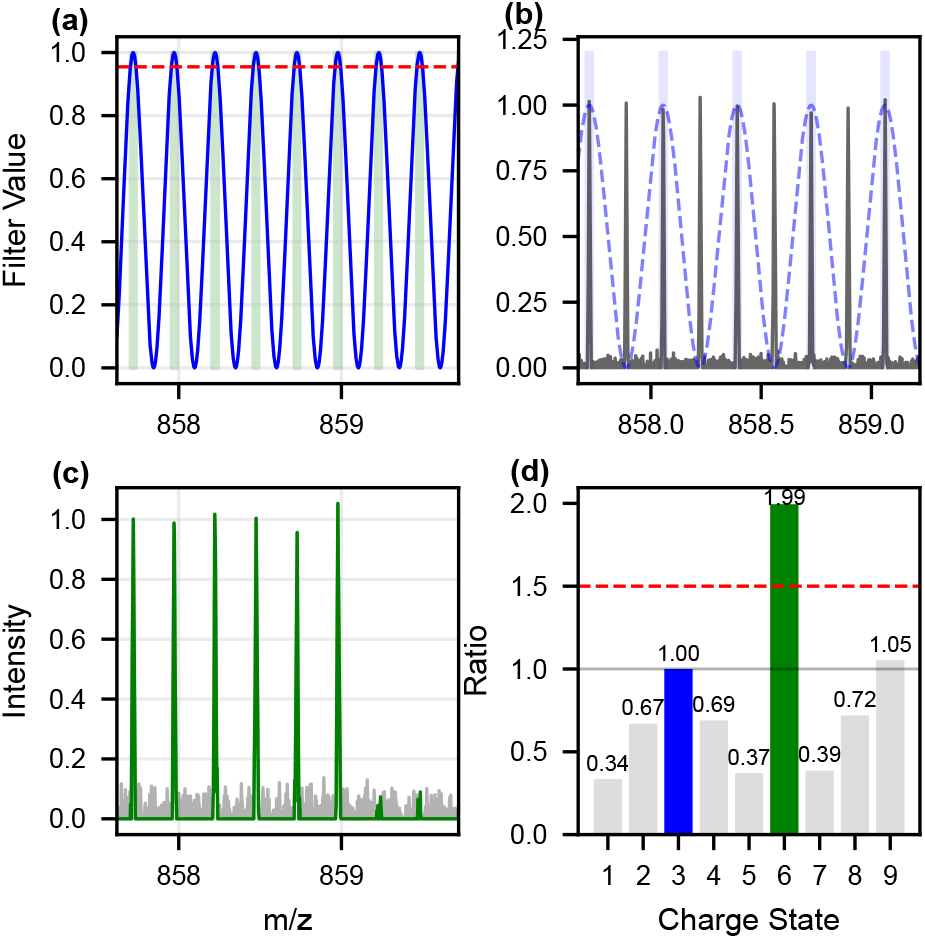
Cosine-squared charge-state filter and aliasing demonstration. **(a)** Filter generation for *z* = 4: the cos^2^ response (blue) with a PPM-derived cutoff (red dashed) defines allowed spectral windows (green shading). **(b)** Aliasing effect: a *z* = 3 filter (blue dashed, blue shading) applied to a true *z* = 6 signal (black) captures every other peak because of harmonic overlap. **(c)** Filter application on a synthetic *z* = 4 spectrum, showing the raw signal (gray) and filtered output (green). **(d)** Aliasing ratio across candidate charge states relative to *z* = 3; the target charge (blue) and true charge (green) are highlighted. The red dashed line marks the 1.5-fold threshold.

### Quality Control Flagging

Each peptide-level output carries a set of quality control flags. Peptides with the OK flag are considered usable; peptides with one or more flags should be excluded from downstream analysis unless explicitly reviewed. The flags and their triggering conditions are as follows.

#### Bad data

Set if no tractable signal was identified anywhere within the peptide’s search window, when no candidate retention interval passed the per-scan *R*^2^ threshold either on the raw spectra or after moving-average smoothing as described under *Initial Seed Interval*. This indicates that the peptide could not be located in the run, and all downstream outputs for this peptide are absent or null.

#### Bad XIC quality

Set if the deconvolution *R*^2^ of the final, aggregated XIC envelope falls below 0.95. This indicates that the observed envelope cannot be cleanly reconstructed from the peptide’s theoretical all-H distribution across allowed deuteration states, and the deconvolved deuteration distribution should not be trusted.

#### Apex outlier

Set if the tentative apex was rejected by the run-wide deviation check described under *Initial Seed Interval* and no lower-ranked candidate sufficiently close to the expected retention time could be substituted. This indicates that no part of the search window contains a signal whose elution behavior is consistent with the run’s overall chromatography.

#### Higher charge state detected

Set if the cosine-squared aliasing mask described under *Overlapping Charge State Interference: Anti-Aliasing* detects an alias-to-expected intensity ratio of at least 1.5 within the recovered retention interval. This indicates that a higher-charge species overlaps the target envelope at a level that may meaningfully distort the deconvolved deuteration distribution.

#### Short retention time

Set if the recovered retention interval is shorter than the user-specified minimum expected duration, or if its duration is a statistically short outlier relative to the run-wide distribution of interval durations. This indicates that the peptide eluted over too narrow a window to support reliable per-scan deconvolution and aggregation.

#### Broad retention time

Set if the recovered retention interval is longer than two standard deviations above the run-wide mean of expected interval durations. The check applies only to deuterated samples, and is waived for peptides whose own expected duration already exceeds that bound, since these are broad by design rather than by mis-assignment. This indicates that the recovered interval likely extends past the peptide’s true elution, admitting co-eluting signal or background into the aggregated envelope.

#### Apex interval inconsistency

Set if the recovered retention interval overlaps its all-H reference interval by an intersection-over-union of 0.25 or less, after correction for the local retention time offset between the two runs. That offset is the median apex deviation among peptides expected to elute within the same half-minute window, falling back to the run-wide median where fewer than ten such peptides are available. This indicates that the peptide was located at a retention time inconsistent with its all-H counterpart, and that the recovered interval may correspond to a different species.

#### Processing error

Set if the peptide could not be carried through a stage of the pipeline, rather than having failed a quality criterion. This occurs when rewindowing of the search result raises an exception or returns no result, when no candidate retention intervals are available at the start of interval preprocessing, when the expected apex time or expected interval duration is missing or negative, when the run-wide retention time offset cannot be estimated, or when the recovered and reference intervals are degenerate and produce a zero union. This indicates a failure of upstream processing or of the reference metadata rather than a judgement about the peptide’s data, and all downstream outputs for this peptide are absent or null.

## Results and Discussion

### Performance

To evaluate the msengine pipeline’s reproducibility, we used the 181-residue SET1 and 327-residue SWD2 protein complex and assessed chromatographic extraction, deuterated-centroid quantification, and peptide identification across eight LC–MS/MS acquisitions spanning four exchange times and two orders of magnitude, as summarized in Figure 6. Sequences of SET1 and SWD2 were concatenated for this analysis, and thus the proteins are not referred to individually. We also prepared a sample of myoglobin that was 50% undeuterated and 50% fully deuterated by spike-in to demonstrate the ability of msengine to capture the complete elution window of a low signal-to-noise ratio (SNR) ion (Figure 7).

**Figure 6.**
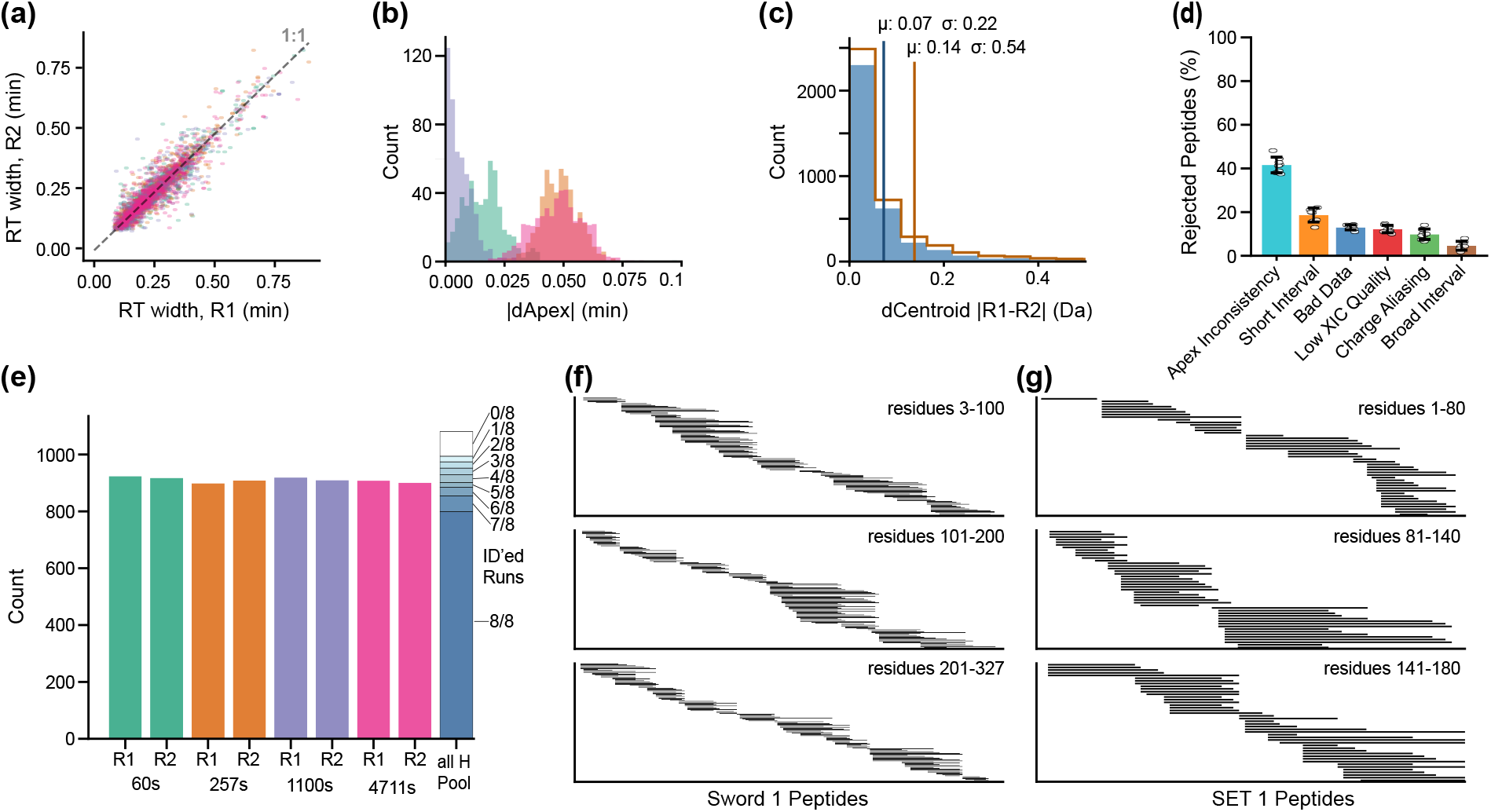
Replicate time-series comparison and peptide coverage. **(a)** Replicate-pair retention-time (RT) window widths. **(b)** Paired apex-time differences. In (a) and (b), colors denote exposure-time pairs: 60 s, green; 257 s, orange; 1100 s, purple; 4711 s, magenta. **(c)** Absolute centroid-difference distributions for accepted peptides (blue) and all non-bad-data matches (orange outline). **(d)** Rejection-flag frequencies across replicates. SET1 and SWD2 panels show peptides accepted in all eight deuterated replicates; outlines indicate detection in the all-H pool. **(e)** Accepted peptide counts per replicate, using the same pair colors; the all-H pool is black, and the final blue stacked bar shows exact deuterated coverage depth.

**Figure 7.**
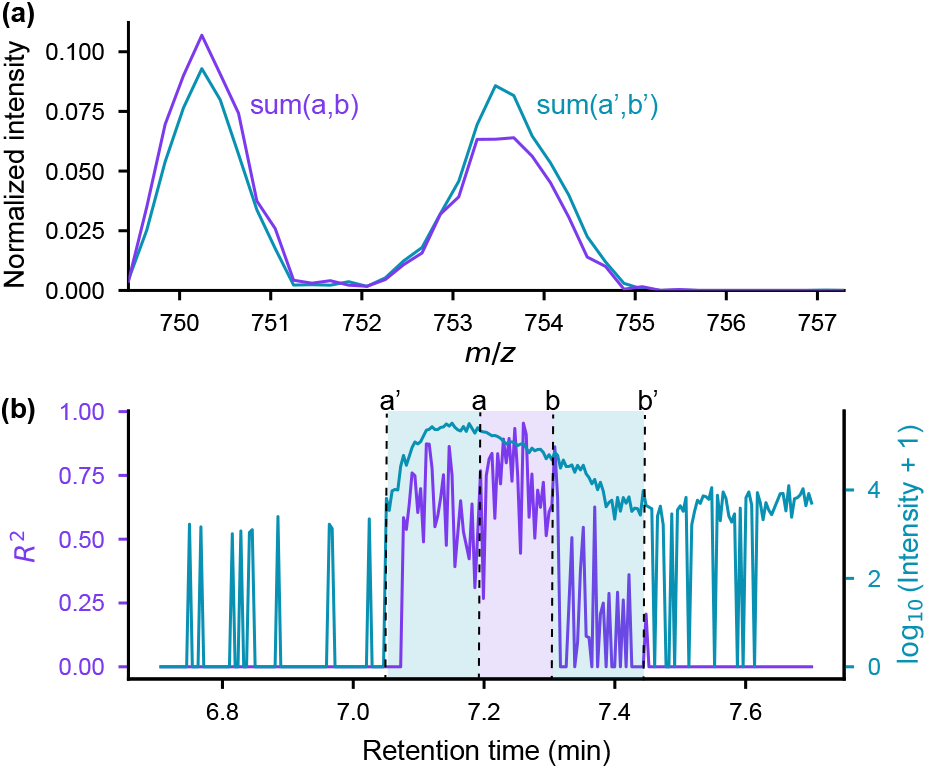
Retention-time selection test to recover a 50:50 mixture for a representative low-SNR peptide. **(a)** Normalized isotopic envelope from the initial *R*^2^-maximizing selection (violet; sum of a and b in panel b) and the final expanded selection (teal; sum of a^*′*^ and b^*′*^ in panel b). The centroid shift between the two selections quantifies the redistribution of isotopic intensity. **(b)** *R*^2^ (violet, left axis) and log-transformed total XIC intensity (teal, right axis) across retention time. Shaded regions indicate the initial *R*^2^-selected interval (violet) and the final expanded interval (teal).

### Peptide Identification

From an initial set of 1,318 possible SET1/SWD2 peptide ions, msengine identified 1,074 (81.5%) that passed all checks in the all-H pool. Of these, 783 (72.9%) were identified in every deuterated measurement, providing greater than 99% sequence coverage and 14-fold redundancy. On average, 901 were found per measurement, representing 83.9% of the all-H pool.

Among the 244 peptides that were present in the input data but not identified in the all-H data, 78 had an outlier apex time, which, if incorrectly assigned in the input list, would represent a false-assignment rate of 5.9%. Another 82 peptides were rejected because of low XIC quality; these peptides were among the lowest-intensity signals in the pool and generally had an SNR below the threshold of 20 needed to pass spectral filtering. Together, these two categories accounted for 65.6% of the peptides not identified in the all-H data.

Three msengine flags that eliminate peptides are expected to occur more frequently as a direct consequence of deuteration: short intervals, inconsistent intervals, and low XIC quality. These issues occur because the spectral envelope broadens, potentially including multiple populations of the same ion, as the peptide exchanges with solvent and takes on additional possible masses. Although we have not computed precise expectation values, deuteration increases the probability that one peptide interferes with another. In such cases, msengine should either shift the extraction window to avoid the interfering ion, producing an interval-inconsistency flag, or truncate the interval, producing a short-interval flag. Although either action may be necessary to avoid contamination, retaining shifted or truncated intervals can inaccurately represent the true chromatographic envelope while still producing an acceptable spectral score. Low XIC quality can arise for a related reason: the signal of any one molecule decreases relative to the total population of masses comprising that peptide as the ensemble becomes distributed over a greater number of possibilities. Thus, a peptide with an SNR above the threshold of 20 in the all-H data may drop below that threshold upon deuteration. On average, 73.6% of rejected peptides were rejected for one of these anticipated issues (Figure 6d). An average of 18.2 peptides (10.6% of rejections) were eliminated because of charge aliasing. The remaining flags were more variable and comprised an average of 24–32 peptides per replicate.

### Extraction and Temporal Precision

Retention-time comparisons between replicates (Figure 6a) demonstrate the adaptive nature of dynamic boundary expansion. Although extraction-window widths (0.08–0.99 min) correlated well (*R*^2^ *≈* 0.88) between replicates, localized variance around the diagonal highlights that boundaries are determined independently per run rather than rigidly matching a specified target width. We observed no change in the expected width with exposure time, indicating that RT windows were not affected by the isotopic broadening that occurs because of progressive deuteration. This fidelity translated to high temporal precision (Figure 6b): median absolute apex-time differences (dApex) ranged from 0.005 to 0.049 min, and all 783 globally shared peptides maintained a per-peptide apex standard deviation of less than 0.1 min across all replicates.

### Benchmarking Performance

A major goal in this work was to match the fidelity of ExMS with fewer heuristics for extracting deuterated mass envelopes and to make better use of computational resources through multithreading and enhanced portability. A comparison of the two programs demonstrates that we achieved this goal.

ExMS identified 488 peptides in every run, with an average of 636 peptides per run, providing 97% coverage with 10-fold redundancy per peptide. Mean absolute apex-time deviations between replicates and centroid errors were marginally larger than those from msengine, indicating that msengine slightly improves replicate extraction precision while significantly increasing the number of peptides identified. Despite processing more peptides than ExMS2, msengine performed better in every reported category and offered performance gains that depended on available hardware. The Computational Performance Benchmarking section of the Supporting Information provides additional performance metrics and a comparison with execution in a Slurm environment.

### Assumption-Free Envelope Verification

Most established approaches for assigning a deuterated envelope to a peptide impose a distributional prior on the extracted signal (binomial or site-probability models, Fourier deconvolution, or model-based deconvolution).^5,7,14–18^ There, the data is fit to an assumed kernel and reads out its parameters. That fit also assigns envelope modality and determines whether the assignment is feasible. The kernel choice therefore biases all three: the extracted quantity is the deuteration distribution’s projection onto the assumed model, and the inferred modality and acceptance of the fit inherit that bias.

In msengine, forward substitution imposes no population shape kernel. The deuterated envelope is reconstructed from the peptide’s all-H isotopic distribution, which is not an assumed shape but the measured basis of every deuterated state. The method asks only whether the observed envelope can be constructed as a non-negative superposition of integer-mass-shifted copies of that basis, and returns the per-population deuterium-uptake vector 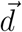 as described under *Envelope Validation*. It introduces no abstraction to modality or binomial form, so the readout is the measured uptake distribution, not the parameters of a model fit to it.

Multimodal envelopes arise through mixed EX1/EX2 exchange and partial cooperative unfolding.^19–23^ A kernel-based readout must either lower its goodness-of-fit thresholds to accept the poor fit, misassign the discrete populations it cannot represent, or compare models containing different numbers of populations. Figure 3 shows an envelope built from four exchange populations, including a sub-population shifted four bins from the main distribution. A kernel-based method treats this shifted intensity as an anomaly; forward substitution reconstructs the full uptake distribution across the envelope from the observed signal alone, without resolving or labeling the underlying populations.

The approach is also far less computationally expensive than iterative model fitting, making scan-by-scan execution practical, and it integrates with noise filtering, localized mass-drift correction, and multipopulation fitting. It provides 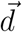 as the direct input for subsequent population deconvolution; kernel assignment and population determination are applied optionally at that stage, as described under *Multipopulation F-Test Plugin*.

### Dynamic Spectral Binning in HDX–MS

Dynamic binning creates isotopologue-aware *m/z* acceptance bins whose widths and centers adapt to the data rather than being fixed in advance. A static scheme centers a fixed-width bin on the monoisotopic peak and shifts it by *k/z* across *n* isotopologue groups, applying the same tolerance at every position. The msengine software instead sizes each bin to the peaks that actually resolve given the ppm error and resolving power of the run, and refines those bins once the deuteration distribution is known.

In deuterated mass spectra, cumulative mass error comprises three distinct factors: uniform offsets from correlated system drift, random offsets from uncorrelated noise, and underlying theoretical error. As instrument resolution improves and random variance diminishes, this theoretical component becomes a primary source of measurement inaccuracy. Because the shape of the deuterated distribution is unknown before measurement, the bins must accommodate every theoretically feasible deuteration state, including the coalescence and overlap between adjacent states; capturing signal this broadly introduces unavoidable center-of-mass shifts. These shifts of the true deuterated-population centers from the static bin centers can be large, as demonstrated in Figure 4.

The pipeline addresses these shifts with the procedure described under *Dynamic ppm Recalibration*. An anomalous peak from an unrelated peptide can fall within the edge of the broad initial bins and corrupt the first deconvolved distribution. That same solution, however, supplies the coefficients used to narrow the bin widths and re-center them on the supported deuterated positions. On re-extraction against these tightened bins, a peak whose position no longer falls within the adjusted tolerance window is excluded, and the distribution is resolved without it. The correction is therefore a self-consistency check: signals that cannot be physically produced by the solved distribution at the given resolving power and ppm tolerance fall outside the refined bins and drop out, rather than being removed by an assumed error model.

### Aliasing

Higher charge states can alias into the spectrum of a lower-charge-state molecule and potentially masquerade as a different ion. For example, a 2+ ion that is isobaric with a different 1+ ion can appear to be the 1+ ion when every other peak is extracted. This problem is related to broader MS work on charge-state assignment, deisotoping, monoisotopic-mass determination, and overlapping isotope distributions.^8–10,24–26^ We address this interference across two distinct layers: the “microstate” of overlapping raw-data curves and the “macrostate” in which valid envelope data are aliased by a higher-frequency, differently charged species.

At the macrostate level, msengine applies the dual-thresholded cosine-squared mask described under *Overlapping Charge State Interference: Anti-Aliasing* to detect when a higher-charge species aliases into the target envelope. On detection, the RT window is re-evaluated, narrowed, or the XIC is flagged. Enumerating all possible interfering peptides from the pool would resolve this for known ions but fails for unidentified ones, and FFT-based detection is too costly to apply scan-by-scan. Our approach is fast enough to run per-scan and feed directly back into RT-window selection.

To resolve microstate overlaps, standard decomposition approaches typically force a strict tradeoff between computationally fast but noise-sensitive algebraic methods and accurate but computationally expensive model-based fitting.^7–10,26^ As outlined under *Multimodal Peak Determination: Shape-Ratio Analysis*, our implementation of Partial Area Gaussian Estimation (PAGE) bridges this gap. By rapidly estimating initial parameters and sequentially evaluating model complexity, PAGE offers flexible resolving power for mixed distributions without requiring the computational burden of traditional fitting.

### Retention-Time Expansion

To ensure comprehensive peak integration, we implemented a dynamic retention-time (RT) selection and expansion algorithm. This approach identifies a reliable core of spectral matches, the seed interval, and extends its boundaries to capture the full chromatographic peak. Rather than requiring each additional scan to pass the seed-stage goodness-of-fit threshold, the expansion terminates on observable chromatographic features: a valley in the intensity trace followed by recovery, or intensity falling below a noise-derived floor for a sustained run of scans. Additionally, if the expanded interval encroaches on adjacent charge-state distributions, the window is subsequently contracted by the anti-aliasing screen to minimize signal overlap. By defining boundaries through these local features, the algorithm incorporates low-intensity scans whose individual fits are weak but whose signal is genuine. This makes measured deuterium-uptake profiles less sensitive to the choice of fit threshold and more representative of the full elution profile.

To demonstrate the ability of msengine to capture full RT windows using its expansion algorithm at the edge of acceptable SNR, and the importance of doing so for measurement of deuterium incorporation, a sample of myoglobin was fully deuterated, mixed 1:1 with a nondeuterated sample, and injected into the mass spectrometer. Every envelope should therefore contain comparable light and heavy populations. Even with an *R*^2^ threshold of 0.7, the true elution window can be obscured at low SNR during scan-by-scan analysis by baseline noise or low intensity that prevents recognition of the mass envelope of interest. For the representative peptide in Figure 7, the seed interval covers less than half of the expanded interval, and restricting extraction to it under-represents the heavy population relative to the light one. Expansion shifts the normalized envelope toward the expected balance. In strongly multimodal measurements, truncating the elution window in this way skews the isotopic envelope, distorts the calculation of deuterium uptake, and can lead to misinterpretation of protein structural dynamics.

### Plugin Compatibility

We designed msengine to be adaptable to users who want a plug-and-play solution for XIC extraction, users who want to adjust outputs through a plugin interface for postprocessing pipelines, and developers who wish to modify the core compute engine directly. It is our intention that the library remain open source and welcoming to contributors of all technical backgrounds. Providing a performant, open-source engine that can extract complete mass envelopes without imposing distributional priors, regardless of instrument resolution or mass error, is important for the accessibility and advancement of the HDX–MS field.

### Multipopulation F-Test Plugin

A foundational design principle of msengine is the deliberate decoupling of raw-signal extraction from distributional modeling. However, most practitioners are interested in population decomposition, so we provide this information natively through a custom postprocessing plugin for multipopulation binomial deconvolution. The plugin also demonstrates how msengine data can be modified through plugin support.

This plugin interfaces directly with the extracted envelope coefficients to generate and fit a composite mathematical model representing overlapping binomial distributions. To prevent statistical overfitting to ambient instrumental noise, the script uses a sequential Ftest framework. It iteratively evaluates the sum of squared errors (SSE) as model complexity increases, terminating the expansion when the addition of a new subpopulation fails to provide a statistically significant improvement (*p ≥* 0.05).

This architecture allows researchers to model complex kinetic behaviors—such as EX1 or mixed-regime exchange dynamics—without the initial extraction algorithm artificially smoothing or distorting discrete structural states. It also highlights the software’s adaptability: using the provided plugin class structure, users can deploy custom physical models, statistical validations, or specialized data transformations directly in the automated post-processing pipeline.

## Conclusions

<monospace>msengine</monospace> extends the retention-time-aware extraction framework established by ExMS and ExMS2 by assigning signal through spectral compatibility rather than conformity to a prescribed deuterium-population distribution. Instead of requiring a binomial, Gaussian, or smoothly evolving envelope, it tests whether each observed spectrum can be reconstructed as a nonnegative superposition of theoretical all-H isotope envelopes shifted to represent integer deuterium counts. A lower-triangular solve makes this criterion practical scan by scan. Resolving-power-aware binning and deuteration-informed refinement restrict extraction to supported spectral regions, while anti-aliasing and quality-control flags identify intervals in which target isolation is unreliable.

Chromatographic expansion complements this spectral criterion by recovering low-intensity scans that may contain different exchange populations. The 1:1 undeuterated/fully deuterated myoglobin mixture illustrates why this matters: selecting scans primarily by spectral fit can truncate the chromatographic profile and distort a multimodal envelope. Chromatographic coverage therefore affects not only signal recovery but also the representation of exchange populations.

In the SET1/SWD2 benchmark, <monospace>msengine</monospace> retained a larger and more persistent accepted target set than ExMS2 while showing lower mean replicate-centroid disagreement. In the absence of an independent extraction-level reference, these comparisons establish differences in accepted-set yield, persistence, and repeatability rather than the correctness of every method-specific extraction. The multiprocessing Python implementation also reduced wall time in the reported same-workstation comparison, at the cost of higher memory use, and supports both workstation and Slurm execution. Together, these developments separate signal extraction from downstream population modeling and help preserve complex exchange behavior as HDX–MS spectra are converted into quantitative measurements.

## Data and Code Availability

The source code for <monospace>msengine</monospace> is maintained at *https://github.com/Genentech/msengine*. Version 0.2.0 was used for the analyses reported here. The raw and processed myoglobin and SET1/SWD2 data have been deposited in MassIVE. The GitHub repository and MassIVE datasets are currently under restricted access and will be made publicly available upon acceptance of the manuscript for peer-reviewed publication. Prior to public release, advance access to the code may be provided upon request to the corresponding author.

## Supporting Information

Supporting Information: PAGE theoretical background, computational-performance bench-marking and benchmark tables, and an experimental-data description (PDF).

## Acknowledgments

The authors thank Genentech for supporting this research. We are also grateful to Mela Mulvihill and Zechariah Philip for proofreading the manuscript and to Aaron Lictao for assistance with data collection. We thank Peter Hsu and Gladys Boenig for providing the SET1/SWD2 protein material used to test the software. During manuscript preparation, the authors used ChatGPT (OpenAI) for editorial review. The authors take full responsibility for the final manuscript.

## References

1. Mayne, L.; Kan, Z.-Y.; Sevugan Chetty, P.; Ricciuti, A.; Walters, B.; Englander, S. Many Overlapping Peptides for Protein Hydrogen Exchange Experiments by the Fragment Separation-Mass Spectrometry Method. Journal of the American Society for Mass Spectrometry 2011, 22, 1898–905.

2. Stofella, M.; Grimaldi, A.; Smit, J. H.; Claesen, J.; Paci, E.; Sobott, F. Computational Tools for Hydrogen–Deuterium Exchange Mass Spectrometry Data Analysis. Chemical Reviews 2024, 124, 12242–12263.

3. Kan, Z.-Y.; Ye, X.; Skinner, J. J.; Mayne, L.; Englander, S. W. ExMS2: An Integrated Solution for Hydrogen–Deuterium Exchange Mass Spectrometry Data Analysis. Analytical Chemistry 2019, 91, 7474–7481.

4. Kan, Z. Y.; Mayne, L.; Chetty, P. S.; Englander, S. W. ExMS: data analysis for HX-MS experiments. J Am Soc Mass Spectrom 2011, 22, 1906–15.

5. Kan, Z. Y.; Walters, B. T.; Mayne, L.; Englander, S. W. Protein hydrogen exchange at residue resolution by proteolytic fragmentation mass spectrometry analysis. Proc Natl Acad Sci U S A 2013, 110, 16438–43.

6. Lindner, R.; Lou, X.; Reinstein, J.; Shoeman, R. L.; Hamprecht, F. A.; Winkler, A. Hexicon 2: Automated Processing of Hydrogen-Deuterium Exchange Mass Spectrometry Data with Improved Deuteration Distribution Estimation. Journal of the American Society for Mass Spectrometry 2014, 25, 1018–1028.

7. Guttman, M.; Weis, D. D.; Engen, J. R.; Lee, K. K. Analysis of Overlapped and Noisy Hydrogen/Deuterium Exchange Mass Spectra. Journal of the American Society for Mass Spectrometry 2013, 24, 1906–1912.

8. Horn, D. M.; Zubarev, R. A.; McLafferty, F. W. Automated Reduction and Interpretation of High Resolution Electrospray Mass Spectra of Large Molecules. Journal of the American Society for Mass Spectrometry 2000, 11, 320–332.

9. Jaitly, N.; Mayampurath, A.; Littlefield, K.; Adkins, J. N.; Anderson, G. A.; Smith, R. D. Decon2LS: An Open-Source Software Package for Automated Processing and Visualization of High Resolution Mass Spectrometry Data. BMC Bioinformatics 2009, 10, 87.

10. Mayampurath, A. M.; Jaitly, N.; Purvine, S. O.; Monroe, M. E.; Auberry, K. J.; Adkins, J. N.; Smith, R. D. DeconMSn: A Software Tool for Accurate Parent Ion Monoisotopic Mass Determination for Tandem Mass Spectra. Bioinformatics 2008, 24, 1021– 1023.

11. Runnalls, A. R. Kullback-Leibler Approach to Gaussian Mixture Reduction. IEEE Transactions on Aerospace and Electronic Systems 2007, 43, 989–999.

12. Zhang, Z.; Smith, D. L. Determination of amide hydrogen exchange by mass spectrometry: A new tool for protein structure elucidation. Protein Science 1993, 2, 522–531.

13. Eibl, G.; Bernardo, K.; Koal, T.; Ramsay, S. L.; Weinberger, K. M.; Graber, A. Isotope correction of mass spectrometry profiles. Rapid Communications in Mass Spectrometry 2008, 22, 2248–2252.

14. Chik, J. K.; Vande Graaf, J. L.; Schriemer, D. C. Quantitating the Statistical Distribution of Deuterium Incorporation To Extend the Utility of H/D Exchange MS Data. Analytical Chemistry 2006, 78, 207–214.

15. Palmblad, M.; Buijs, J.; Håkansson, P. Automatic Analysis of Hydrogen/Deuterium Exchange Mass Spectra of Peptides and Proteins Using Calculations of Isotopic Distributions. Journal of the American Society for Mass Spectrometry 2001, 12, 1153–1162.

16. Hotchko, M.; Anand, G. S.; Komives, E. A.; Ten Eyck, L. F. Automated Extraction of Backbone Deuteration Levels from Amide H/2H Mass Spectrometry Experiments. Protein Science 2006, 15, 583–601.

17. Na, S.; Lee, J.-J.; Joo, J. W. J.; Lee, K.-J.; Paek, E. deMix: Decoding Deuterated Distributions from Heterogeneous Protein States via HDX-MS. Scientific Reports 2019, 9, 3176.

18. Saltzberg, D. J.; Broughton, H. B.; Pellarin, R.; Chalmers, M. J.; Espada, A.; Dodge, J. A.; Pascal, B. D.; Griffin, P. R.; Humblet, C.; Sali, A. A Residue-Resolved Bayesian Approach to Quantitative Interpretation of Hydrogen-Deuterium Exchange from Mass Spectrometry: Application to Characterizing Protein-Ligand Interactions. The Journal of Physical Chemistry B 2017, 121, 3493–3501.

19. Weis, D. D.; Wales, T. E.; Engen, J. R.; Hotchko, M.; Ten Eyck, L. F. Identification and Characterization of EX1 Kinetics in H/D Exchange Mass Spectrometry by Peak Width Analysis. Journal of the American Society for Mass Spectrometry 2006, 17, 1498–1509.

20. Ferraro, D. M.; Lazo, N. D.; Robertson, A. D. EX1 Hydrogen Exchange and Protein Folding. Biochemistry 2004, 43, 587–594.

21. Engen, J. R.; Wales, T. E.; Chen, S.; Marzluff, E. M.; Hassell, K. M.; Weis, D. D.; Smithgall, T. E. Partial Cooperative Unfolding in Proteins as Observed by Hydrogen Exchange Mass Spectrometry. International Reviews in Physical Chemistry 2013, 32, 96–127.

22. Tuttle, L. M.; James, E. I.; Georgescauld, F.; Wales, T. E.; Weis, D. D.; Engen, J. R.; Nath, A.; Klevit, R. E.; Guttman, M. Rigorous Analysis of Multimodal HDX-MS Spectra. Journal of the American Society for Mass Spectrometry 2025, 36, 416–423.

23. Fang, J.; Rand, K. D.; Beuning, P. J.; Engen, J. R. False EX1 Signatures Caused by Sample Carryover During HX MS Analyses. International Journal of Mass Spectrometry 2011, 302, 19–25.

24. Senko, M. W.; Beu, S. C.; McLafferty, F. W. Automated Assignment of Charge States from Resolved Isotopic Peaks for Multiply Charged Ions. Journal of the American Society for Mass Spectrometry 1995, 6, 52–56.

25. Senko, M. W.; Beu, S. C.; McLafferty, F. W. Determination of Monoisotopic Masses and Ion Populations for Large Biomolecules from Resolved Isotopic Distributions. Journal of the American Society for Mass Spectrometry 1995, 6, 229–233.

26. Hoopmann, M. R.; Finney, G. L.; MacCoss, M. J. High-Speed Data Reduction, Feature Detection, and MS/MS Spectrum Quality Assessment of Shotgun Proteomics Data Sets Using High-Resolution Mass Spectrometry. Analytical Chemistry 2007, 79, 5620–5632.

