## Supplementary Information for "msengine: Adaptive Open-Source Signal Extraction for HDX MS"

#### S1 Partial Area Gaussian Estimation (PAGE)

The accurate measurement of peak intensity is often complicated by the overlap of isotopic peaks from molecules with similar mass-to-charge ratios. This overlap can cause significant deviations in both intensity and the weighted center of a peak. Although knowing the exact masses of all nonisomeric molecules involved can permit more rigorous peak-extraction methods, such data are rarely available at the time of centroiding. Consequently, standard practices—such as thresholding, weighted means, apex detection, three-point Gaussian fitting, and the method of moments—often force a trade-off among accuracy, sensitivity to noise, and computational cost.

To address this problem, we developed a shape-ratio analysis approach. The method parameterizes a single Gaussian using the partial area and width around a peak's highest-intensity point. A greedy insertion strategy iteratively adds Gaussians, and F-testing prevents overfitting. This approach reveals peak information missed by apex detection and separates centroids with accuracy comparable to weighted averaging around the apex. The implementation offers a computational middle ground: it is more robust than basic centroiding techniques but less computationally expensive than traditional Gaussian fitting.

##### S1.1 Theoretical Background

We begin by integrating a noisy Gaussian centered at zero.

$$g(x) = Ae^{-x^2/C} + \epsilon, \qquad C = 2\sigma^2 \tag{S1}$$

Here, $\epsilon$ is a random variable with a probability-density function symmetric about a known or calculable mean $\mu$.

$$P_{\text{area}} = 2\int_0^b g(x)\,\mathrm{d}x = A\sqrt{\pi C}\,\operatorname{erf}\left(\frac{b}{\sqrt{C}}\right) + 2b\mu, \tag{S2}$$

$$\frac{P_{\text{area}} - 2b\mu}{A\sqrt{\pi}} = \sqrt{C}\,\operatorname{erf}\left(\frac{b}{\sqrt{C}}\right) \tag{S3}$$

Solving for $\sigma$ analytically is not possible in this form. However, if we define a new variable $\phi$ that relates the known value $b$ to $\sigma$, we can solve the equation numerically.

$$\phi = \frac{b}{\sqrt{C}}, \tag{S4}$$

$$F = \frac{P_{\text{area}}}{bA\sqrt{\pi}} - \frac{2\mu}{A\sqrt{\pi}} = \frac{\operatorname{erf}(\phi)}{\phi} \tag{S5}$$

This expression enables us to solve for $\phi$ using Newton's method when $F$ is known. Once $\phi$ is obtained, $\sigma$ follows from

$$\sigma = \frac{b}{\phi\sqrt{2}} \tag{S6}$$

The algorithm estimates Gaussian parameters by calculating an F-value from the ratio of integrated area to peak amplitude, then inverting the relationship $F = \operatorname{erf}(\phi)/\phi$ to find $\phi$, from which $\sigma$ is extracted. The theoretical formula assumes symmetric integration bounds around the peak center, but this may not always be possible when data is discrete. When the Gaussian is off-center within the data range, the asymmetric bounds cause the calculated F-value to differ from the theoretical $\operatorname{erf}(\phi)/\phi$ relationship. This discrepancy grows with $\sigma$ because wider Gaussians have more area in their tails. This behavior is demonstrated in Figure S3a. Additionally, implicit error arises from the discrete area approximation under the curve, as shown in Figure S3b.

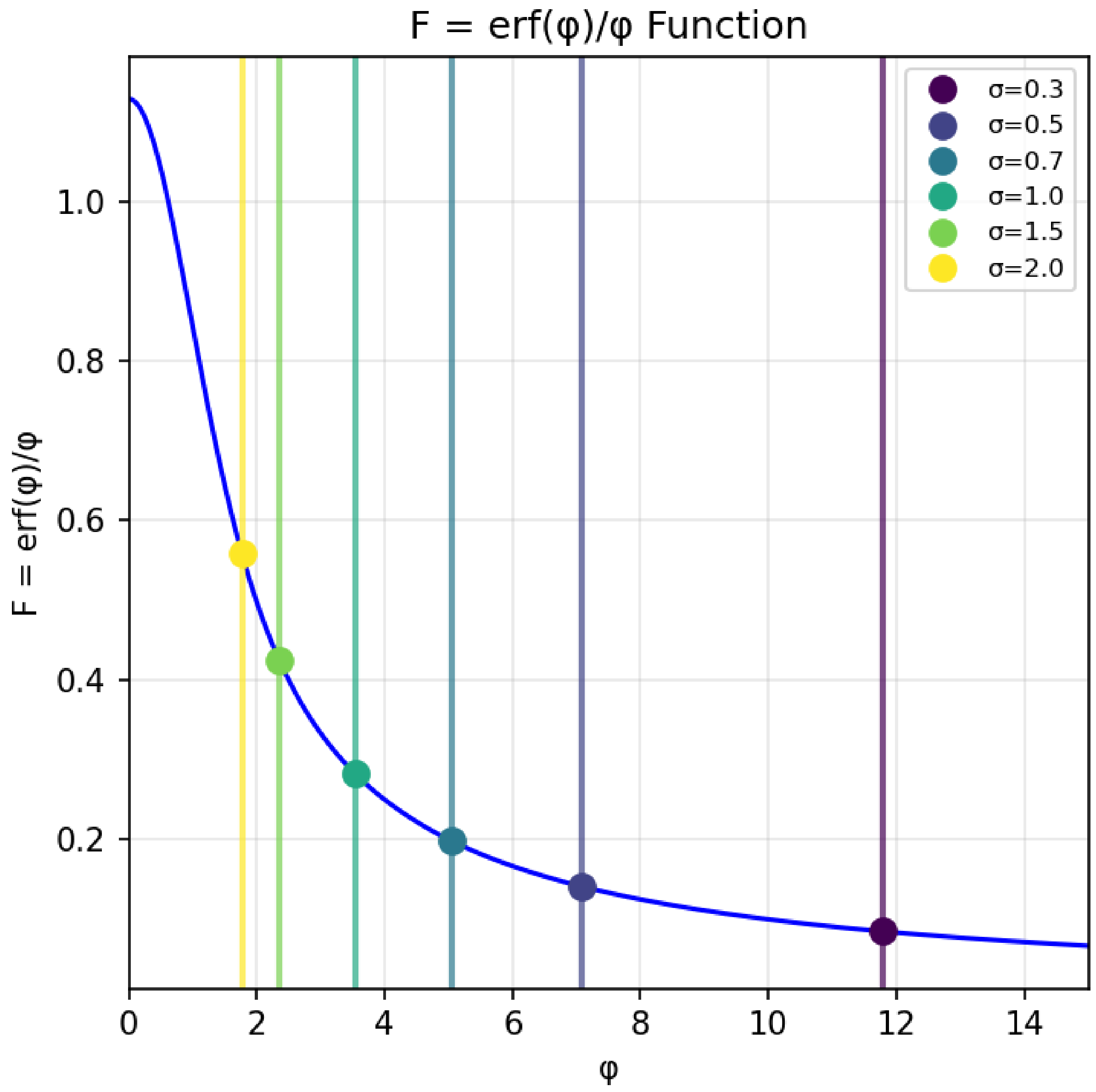


Figure S1: Relationship between $F = \mathrm{erf}(\phi)/\phi$ and $\phi$ used for PAGE width estimation. Colored markers and vertical guides identify the plotted example $\sigma$ values.

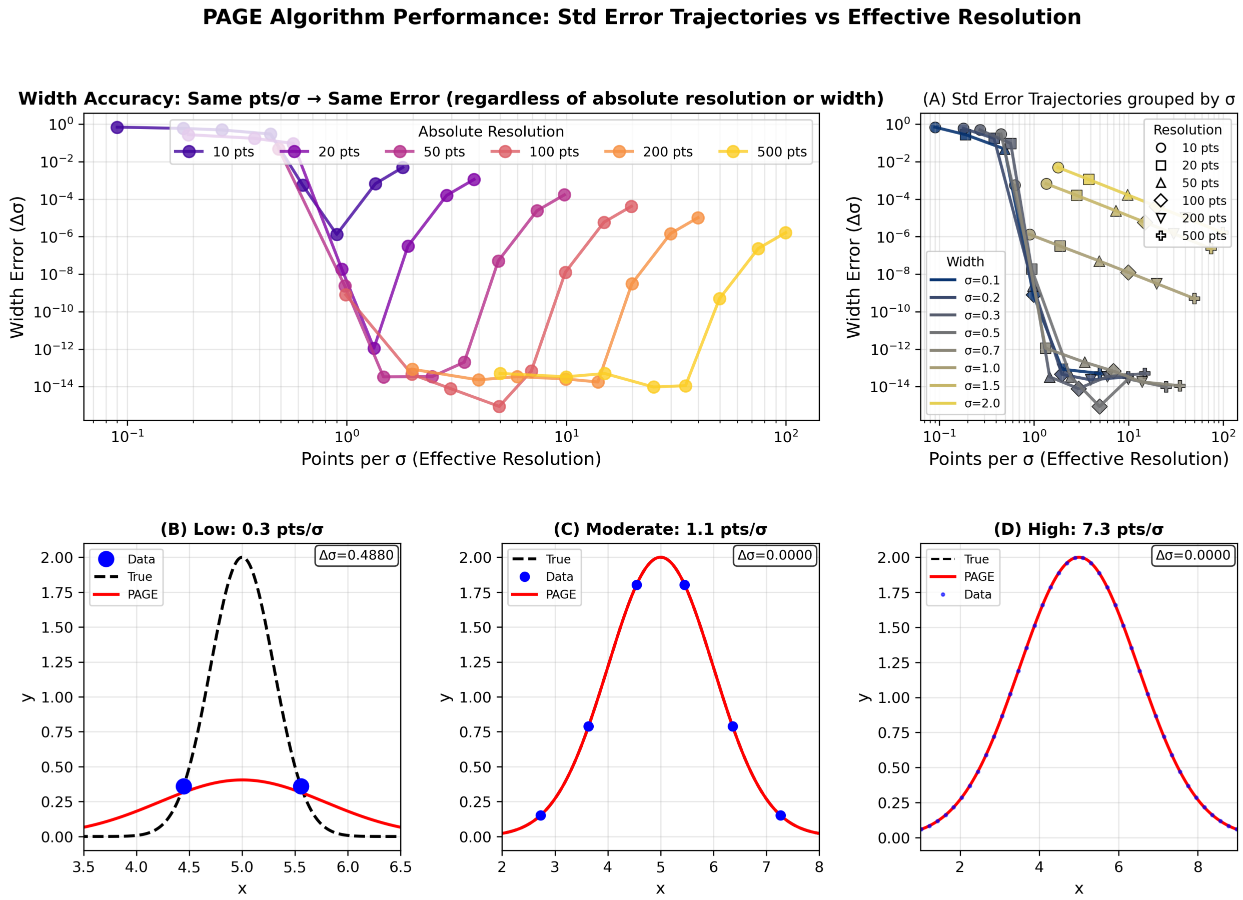


Figure S2: (Top) Width estimation error ($\Delta\sigma$) as a function of effective resolution (points per $\sigma$) for different absolute resolutions (10–500 points). Initial curves collapse onto a single trend, then deviate at higher effective resolutions. This demonstrates the systematic error dependencies shown in Figure S3. (Bottom) Example Gaussian fits at low (0.3 pts/$\sigma$), moderate (1.1 pts/$\sigma$), and high (7.4 pts/$\sigma$) effective resolutions, showing the PAGE estimate (red) compared with the true Gaussian (black dashed) and sampled data points (blue).

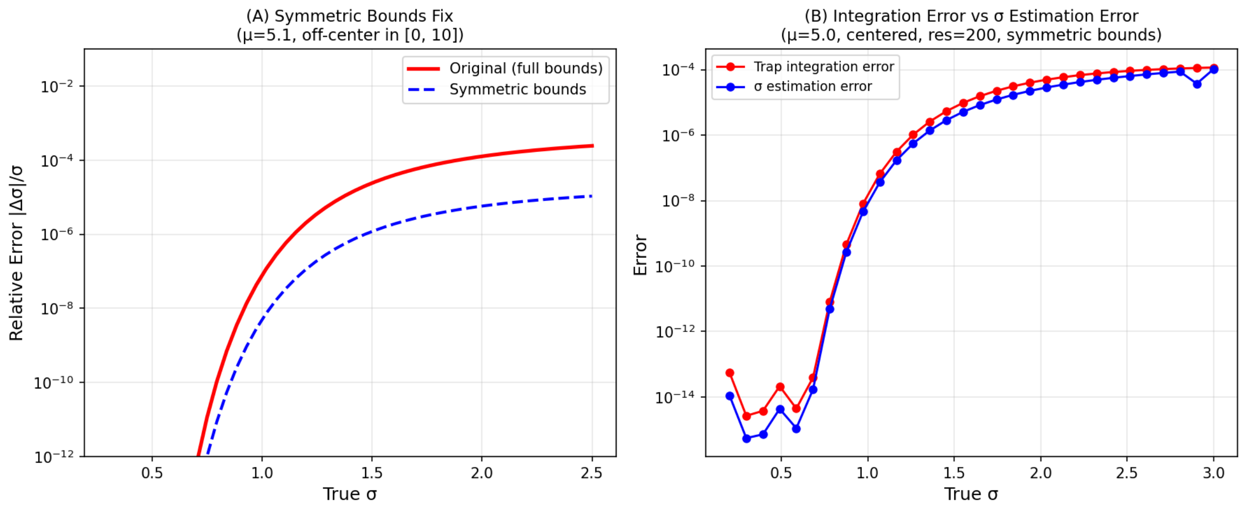


Figure S3: Relative error in Gaussian width ( $\sigma$) estimation as a function of true $\sigma$ for an off-center Gaussian ($\mu = 5.17$) in the domain [0, 10]. The original PAGE algorithm using full integration bounds (red solid line) shows increasing error with Gaussian width, reaching $10^{-4}$ at $\sigma = 2.5$. The reselected optimized bounds for symmetric center on the detected peak (blue dashed line) maintains errors below $10^{-3}$ across all tested widths. (B) Shows error from area integration scheme to be the dominating error for higher effective resolutions. Data generated at 500-point resolution with no added noise.

### S2 Computational Performance Benchmarking

On the same Windows workstation, msengine completed all-H processing in 4.57 min versus 35.06 min for ExMS2 (7.68-fold faster) and deuterated-data processing in 2.76 min versus 36.81 min (13.36-fold faster). Total stage time was 7.32 min for msengine and 71.88 min for ExMS2, a 9.82-fold speedup and an 89.81% reduction in wall time. The msengine runs averaged 12.40 and 10.57 active logical cores during all-H and D processing, compared with 1.06 and 1.00 for ExMS2. Peak process-tree RSS was correspondingly higher for msengine: 13.76 versus 1.79 GiB for all-H and 9.03 versus 2.14 GiB for D.

#### S2.1 Methods

The Windows system was an HP Z4 G4 workstation running 64-bit Windows 10 Enterprise LTSC, with an Intel Core i9-10900X processor (10 physical cores, 20 logical processors; 3.70 GHz), 63.8 GiB RAM, NVMe storage, and the High performance power plan. The msengine version 0.2.0 used Python 3.11.13, 20 multiprocessing workers, one numerical-library thread per worker, a 51.037-GiB memory budget, 5-ppm mass tolerance, 0.05-min minimum RT width, and an $R^2$ threshold of 0.8. D processing additionally used five-population fitting and Xavier export. ExMS2 release 20181103 used its defaults: 10-ppm $m/z$ tolerance, 2-min all-H and 0.5-min D RT windows, individual-peak noise threshold 500, and summed-peak noise threshold 1,500.

Both programs received the same Byonic-filtered `READY_250122_SWD_Set_MSMS.csv` file containing 1,318 distinct peptide–charge candidates (781 unique peptide sequences) that had been already vetted using the Byonic program. The same 1.531-GiB all-H and 1.778-GiB D mzXML files were used for both programs. ExMS2 timing included launch, GUI setup, processing, and user confirmation; The msengine timing covered command launch through process-tree termination. Windows resources were sampled every 0.5 s. Mean active cores equaled cumulative CPU seconds divided by wall seconds, and CPU efficiency divided this value by 20 logical processors. Peak RSS was summed across the process tree.

Table S1: End-to-end performance on the same Windows workstation. Speedup is ExMS2 wall time divided by msengine wall time; each condition was measured once.

| Stage | Wall (min) | | Speedup | CPU (min) | | Peak RSS (GiB) | | Active cores |
|---|---|---|---|---|---|---|---|---|
| | ExMS2 | msengine | | ExMS2 | msengine | ExMS2 | msengine | msengine |
| all-H | 35.06 | 4.57 | 7.68 | 37.12 | 56.65 | 1.79 | 13.76 | 12.40 |
| D | 36.81 | 2.76 | 13.36 | 36.85 | 29.12 | 2.14 | 9.03 | 10.57 |

**Candidate and retained-output normalization:** The primary normalization divided run-level resources by the common 1,318 submitted peptide–charge candidates in order to compare computational load for the same supplied search space, not resources per peptide found. Wall time per candidate is inverse aggregate throughput under parallel execution, and peak RSS per candidate is descriptive rather than marginal memory cost.

Table S2: Run-level resources normalized by the 1,318 submitted peptide–charge candidates (781 unique sequences). Wall time per candidate is inverse aggregate throughput; peak RSS per candidate is descriptive.

| Program | Environment | Stage | Wall (s/candidate) | CPU (s/candidate) | Candidates/s | Peak RSS/candidate (MiB) |
|---|---|---|---|---|---|---|
| ExMS2 | Windows | all-H | 1.596 | 1.690 | 0.626 | 1.39 |
| msengine | Windows | all-H | 0.208 | 2.579 | 4.810 | 10.69 |
| msengine | HPC/Slurm | all-H | 0.244 | 1.372 | 4.093 | 5.61 |
| ExMS2 | Windows | D | 1.676 | 1.678 | 0.597 | 1.66 |
| msengine | Windows | D | 0.125 | 1.326 | 7.972 | 7.01 |
| msengine | HPC/Slurm | D | 0.147 | 0.768 | 6.794 | 5.98 |

ExMS2 retained 1,185 peptides for all-H and 651 for the deuterated dataset, while msengine retained 1,141 peptides for all-H and 998 for D (defined as those passing all skip tests or rows with `Status=OK`, respectively). A secondary normalization divided resources by these program-specific retained counts. Notably, this comparison is not truly quantitative because ExMS2 skip tests and msengine status rules are not equivalent. However, both approaches attempt to eliminate peptides that do not pass quality thresholds, and so the estimates are representative of what each program would deliver on these data.

Table S3: Program-reported outcomes and resources normalized by retained targets. ExMS2 retention required passing all skip tests; msengine retention required `Status=OK`. The quality systems are not equivalent.

| Program | Stage | Analyzed | Found | Retained | Retained (%) | Wall (s/retained) | CPU (s/retained) | Peak RSS/retained (MiB) |
|---|---|---|---|---|---|---|---|---|
| ExMS2 | all-H | 1,309 | 1,276 | 1,185 | 90.5 | 1.775 | 1.880 | 1.55 |
| msengine | all-H | 1,318 | – | 1,141 | 86.6 | 0.240 | 2.979 | 12.35 |
| ExMS2 | D | 1,309 | 1,242 | 651 | 49.7 | 3.393 | 3.397 | 3.37 |
| msengine | D | 1,141 | – | 998 | 87.5 | 0.166 | 1.751 | 9.26 |

**HPC execution:** The msengine software was also run through Slurm using the same inputs. Each dependent all-H and D job used 10 CPUs, `-MP 10`, one numerical-library thread per worker, and 20 GiB allocated memory. Queue time was excluded. All-H took 5.37 min and D took 3.23 min (8.60 min summed execution; 8.77 min dependency- controlled makespan). Individual D jobs can be submitted with additional resources or in parallel, depending only on the first all-H job. These conditions were chosen to reflect equivalence with the Windows machine.

Table S4: msengine Slurm execution. Wall time excludes queue waiting; each job used 10 CPUs and 20 GiB allocated memory.

| Stage | Wall (min) | CPU (min) | Active cores | CPU efficiency (%) | Peak RSS (GiB) | Throughput (MiB/s) |
|---|---|---|---|---|---|---|
| all-H | 5.37 | 30.13 | 5.61 | 56.15 | 7.22 | 4.870 |
| D | 3.23 | 16.88 | 5.22 | 52.21 | 7.69 | 9.386 |

#### S3 Experimental Data

The SET1/SWD2 protein complex was used to assess msengine and ExMS2, provided by colleagues Peter Hsu and Gladys Boenig for testing the program. Myoglobin from equine skeletal muscle was purchased from Sigma as a lyophilized powder. Tandem mass spectra were collected to identify peptides for subsequent tracking during deuterium-exchange experiments and were processed using Byonic to generate the peptide input list.

Deuterated spectra were collected at 60, 257, 1100, and 4711 s in duplicate by diluting 3 $\mu$L of 25 $\mu$M SET1/SWD2 complex with 60 $\mu$L of deuterated 20 mM HEPES, 150 mM NaCl, pH 7.5, using a Trajan DHR HDX robotic system (Trajan Scientific America). After specified labeling times, 55 $\mu$L of this mixture was then diluted 1:1 with quench buffer containing 4 M GdmCl and 1 M glycine at pH 2.5 and then 95 $\mu$L immediately injected into a cold online LC–MS system. Peptides were digested using a Nepenthesin-II column (Affipro), washed for three minutes on a C-18 Vanguard trap column, and separated on a C18 column (Acquity UPLC BEH C18, 1.7 $\mu$m, 2.1 mm × 50 mm, Waters) using a linear gradient of 5–40% acetonitrile in water with 0.1% formic acid over 15 min at a flow rate of 35 $\mu$L/min. Deuterated mass spectra were acquired on a Thermo Scientific Exploris 480 mass spectrometer in positive ion mode with a resolution of 120,000 at m/z 200. Tandem-MS runs for peptide identification uses a top-7 DDA method with a fragment resolution of 15,000 at m/z 200, and parent resolution of 60,000 at m/z 200. Peptide identifications were filtered to a 1% FDR using the Byonic score and reverse-decoy strategy. The resulting peptide list was used as input for both msengine and ExMS2. The SET1/SWD2 raw and processed data have been deposited in MassIVE.

In order to measure a 1:1 undeuterated/perdeuterated myoglobin mixture, pH 7.5 solutions of roughly 0.5 $\mu$M myoglobin were prepared in both $H_2O$ and $D_2O$ and incubated at 30 °C for 24 h. The samples were concentration matched by absorption at 280 nm after incubation. Using the same robotics platform as our SET1/SWD2 measurements, myoglobin mass spectra were acquired by first combining 30 $\mu$L of each sample, followed by a 1:1 dilution with quench buffer before injection onto the same system described above and identical treatment. These samples were measured in triplicate. The raw and processed myoglobin data and peptide lists have been deposited in MassIVE.

The MassIVE datasets are currently under restricted access and will be made publicly available together with the `msengine` source code upon acceptance of the manuscript for peer-reviewed publication.
